# Impact of glycoengineering and immunogenicity on the anti-cancer activity of a plant-made lectin-Fc fusion protein

**DOI:** 10.1101/2022.05.31.494188

**Authors:** Matthew Dent, Katarina L. Mayer, Noel Verjan Garcia, Haixun Guo, Hiroyuki Kajiura, Kazuhito Fujiyama, Nobuyuki Matoba

## Abstract

Plants are an efficient production platform for manufacturing glycoengineered monoclonal antibodies and antibody-like molecules. Avaren-Fc (AvFc) is a lectin-Fc fusion protein or lectibody produced in *Nicotiana benthamiana,* which selectively recognizes cancer-associated high-mannose glycans. In this study, we report the generation of a glycovariant of AvFc that is devoid of plant glycans, including the core α1,3-fucose and β1,2-xylose residues. The successful removal of these glycans was confirmed by glycan analysis using HPLC. This variant, AvFc^ΔXF^, has significantly higher affinity for Fc gamma receptors and induces higher levels of luciferase expression in an antibody-dependent cell-mediated cytotoxicity (ADCC) reporter assay against B16F10 murine melanoma cells without inducing apoptosis or inhibiting proliferation. In the B16F10 flank tumor mouse model, we found that systemic administration of AvFc^ΔXF^, but not an aglycosylated AvFc variant lacking affinity for Fc receptors, significantly delayed the growth of tumors, suggesting that Fc-mediated effector functions were integral. AvFc^ΔXF^ treatment also significantly reduced lung metastasis of B16F10 upon intravenous challenge whereas a sugar-binding-deficient mutant failed to show efficacy. Lastly, we determined the impact of anti-drug antibodies (ADAs) on drug activity *in vivo* by pretreating animals with AvFc^ΔXF^ before implanting tumors. Despite a significant ADA response induced by the pretreatment, we found that the activity of AvFc^ΔXF^ was unaffected by the presence of these antibodies. These results demonstrate that glycoengineering is a powerful strategy to enhance AvFc’s antitumor activity.

## INTRODUCTION

Cancer immunotherapy with monoclonal antibodies (mAbs) targeting tumor-associated antigens (TAAs) has forever altered treatment paradigms and has vastly improved patient survival and quality of life. MAbs exert their anti-cancer activities through a combination of immune-mediated and non-immune-mediated mechanisms such as direct receptor inhibition, antibody-dependent cell-mediated cytotoxicity (ADCC), antibody-mediated phagocytosis, and complement-mediated cell lysis. The initiation of inflammatory responses by antibodies is largely dependent on the binding and activation of Fcγ receptors (FcγRs), which are differentially expressed in several immune cell types, in particular natural killer (NK) cells, neutrophils, macrophages, and monocytes (Mancardi and Daëron, 2014). Binding of the Fc region of an antibody to the activating FcγRs (FcγRI, FcγRIIa, FcγRIIIa) results in the generation of signaling cascades through intracellular ITAM domains, leading to cellular activation, degranulation, or phagocytosis in a cell type-specific manner (Vidarsson et al., 2014). Initiation of ADCC, for instance, is accomplished primarily by recognition of antibody-opsonized cells by FcγRIIIa on NK cells, which subsequently release cytotoxic granules containing granzyme and perforin to initiate target cell death and begin to express IFNγ. As these immune-mediated mechanisms play an important role in the effects of therapeutic mAb drugs, even those whose primary mechanism is receptor antagonization (Weiner, 2010), much research has been conducted into enhancing their ability to activate Fc functions by improving their affinity to the various FcγRs with the goal of improving clinical outcomes (Zahavi et al., 2018).

The strength of the Fc-mediated response reflects both the density of the target TAAs on the cell surface as well as the affinity of the mAb to the FcγR (Ochoa et al., 2017). The affinity of this interaction is determined by both the IgG isotype of the antibody as well as the composition of its N-glycans attached to the Asn297 within the Fc region (Jennewein and Alter, 2017; Vidarsson et al., 2014). One method for improving the affinity of an Fc to the FcγRs is through point mutation of the Fc region. A well-known example of this is the GASDALIE mutation, which is a series of 4 amino acid substitutions in the C_H_2 and C_H_3 domains that significantly increases the affinity of the Fc for FcγRIIIa (Ahmed et al., 2016; Smith et al., 2012). This mutation has been trialed on a number of mAb therapeutics and has consistently resulted in increased ADCC activity and *in vivo* efficacy in pre-clinical models of both viral diseases and cancer (DiLillo and Ravetch, 2015; Edwards et al., 2021). Host glycoengineering is another method commonly used to modify FcγR affinity. N-glycosylation of mAbs occurs at a single conserved site on the C_H_2 domain of the Fc region, the composition of which can be altered through manipulation of host glycosyltransferase enzyme expression (Wang et al., 2019b). This can be performed chemically through exposure to compounds such as kifunensine, which inhibits mannosidase I and results in an abundance of high-mannose-type glycans, or genetically through knockdowns or knockouts of glycosyltransferase genes by RNAi or CRISPR-Cas9 (Hanania et al., 2017; Kommineni et al., 2019; Roychowdhury et al., 2018; Strasser et al., 2008). In plants, recombinant expression of antibodies results in the majority of glycoforms containing β1,2-xylose and α1,3-fucose residues (Schähs et al., 2007). Expression in glycoengineered *Nicotiana benthamiana* containing RNAi knockdowns of β1,2-xylosyltransferase and α1,3-fucosyltransferase genes (ΔXylT/ΔFucT, or ΔXF), however, results in a near total loss of plant-typical glycans with the predominant glycoform consisting of the core trimannose and two N-acetylglucosamine residues (Strasser et al., 2008). Antibodies manufactured in these plants have significantly higher FcγR affinity, which is similar to the effect achieved following removal of the core α1,6-fucose residue from antibodies expressed in mammalian cells (Schähs et al., 2007; Shields et al., 2002). Thus, removal of plant-derived glycans is an attractive target to achieve for therapeutic antibody development, and indeed a number of stable transgenic *N. benthamiana* lines for recombinant protein expression have been generated with that goal in mind (Hanania et al., 2017; Jansing et al., 2019; Strasser et al., 2008; Weterings and Van Eldik, 2019).

Plants, in particular *N. benthamiana,* are quite amenable to glycoengineering owing to the relatively lower complexity of the *N*-glycosylation pathway compared to that of mammalian culture systems, which are currently the industry standard for mAb manufacturing. This makes them an attractive alternative platform that has already been used to manufacture dozens of antiviral and anticancer mAb and Fc-fusion protein therapeutics with glycan or amino acid modifications (Forthal et al., 2010; Hurtado et al., 2020; Lai et al., 2014; Loos et al., 2014; Marusic et al., 2018; Tsekoa et al., 2016). Previously, we have described the development in plants of a novel lectin-Fc fusion protein, or lectibody, which targets cancer-associated high-mannose glycans called Avaren-Fc (AvFc) (Hamorsky et al., 2019; Oh et al., 2021). The presence of aberrant glycosylation patterns on cell surface glycoproteins has been identified as a hallmark of cancer, and an overabundance of high-mannose glycans has been found in numerous human cancers including colorectal cancer (Boyaval et al., 2021; Chik et al., 2014; Sethi et al., 2016), hepatocellular carcinoma (Powers et al., 2015; Takayama et al., 2020), cholangiocarcinoma (Park et al., 2020), lung adenocarcinoma (Ruhaak et al., 2015), pancreatic cancer (Park et al., 2015), ovarian cancer (Chen et al., 2017; Everest-Dass et al., 2016), prostate cancer (Munkley et al., 2016), and some skin cancers (Möginger et al., 2018). This suggests that display of these immature glycans may be common in cancer due to an inherent property of the transformation to malignancy, and this fact can potentially be exploited to create a new druggable target for therapy. We have previously reported that AvFc recognizes a large number of cancer cell lines through this mechanism, and that by binding to mannosylated forms of EGFR and IGF1R derived from lung cancer cell lines and tissues in addition to inducing ADCC, AvFc displays potent activity against A549 and H460 lung cancer both *in vitro* and *in vivo* (Oh et al., 2021). To further explore and examine the contribution of ADCC to AvFc’s antitumor mechanism of action, we have generated a variant of AvFc by expression in ΔXF plants (AvFc^ΔXF^) that is devoid of plant-derived glycans and may hence exhibit greater ADCC activity due to the lack of core fucosylation. In this study, we set out to characterize this variant as well as investigate its activity by comparing it to an aglycosylated variant, AvFc^Δgly^, and a variant lacking sugar-binding activity, AvFc^Δlec^, using both *in vitro* assays as well as the syngeneic murine B16F10 melanoma model. Additionally, we explored the impact of the generation of anti-drug antibodies (ADAs) on the efficacy of AvFc in this model. The results demonstrate the importance of Fc modification on the therapeutic efficacy of AvFc, as well as the utility of the plant expression system for manufacturing glycoengineered AvFc variants.

## RESULTS

### AvFc variants are highly expressed and easily purified in plant tissues

We have previously reported that AvFc was highly expressed in wild-type *N. benthamiana* (AvFc^WT^) by the magnICON tobamovirus vector transient overexpression system and had a purified yield of ≈ 100 mg from one kg of leaf biomass (Hamorsky et al., 2019). Here, AvFc^WT^, AvFc^ΔXF^ and AvFc^Δlec^ were produced using the same manufacturing method. Yields of these purified proteins were not found to be significantly different, averaging between 100 and 150 mg/kg depending on plant conditions at ≈ 95% purity as determined by densitometry analysis of a Coomassie-stained gel (Figure 1A). On the other hand, removal of the single N-glycan in the AvFc^Δgly^ variant resulted in a more than 50% decrease in yield, likely due to a decrease in stability *in planta.* All the variants were indistinguishable according to molecular weight (≈ 38.6 kDa reduced, ≈ 77.1 kDa non-reduced, Figure 1A), though the amounts of a frequently observed 50 kDa band, likely corresponding to Fc dimer fragments, varied somewhat between variants. Thus, we concluded that change in the plant expression host from wild type to the ΔXylT/ΔFucT transgenic line did not significantly alter the manufacturability or the resulting protein purity of AvFc.

**Figure 1.**
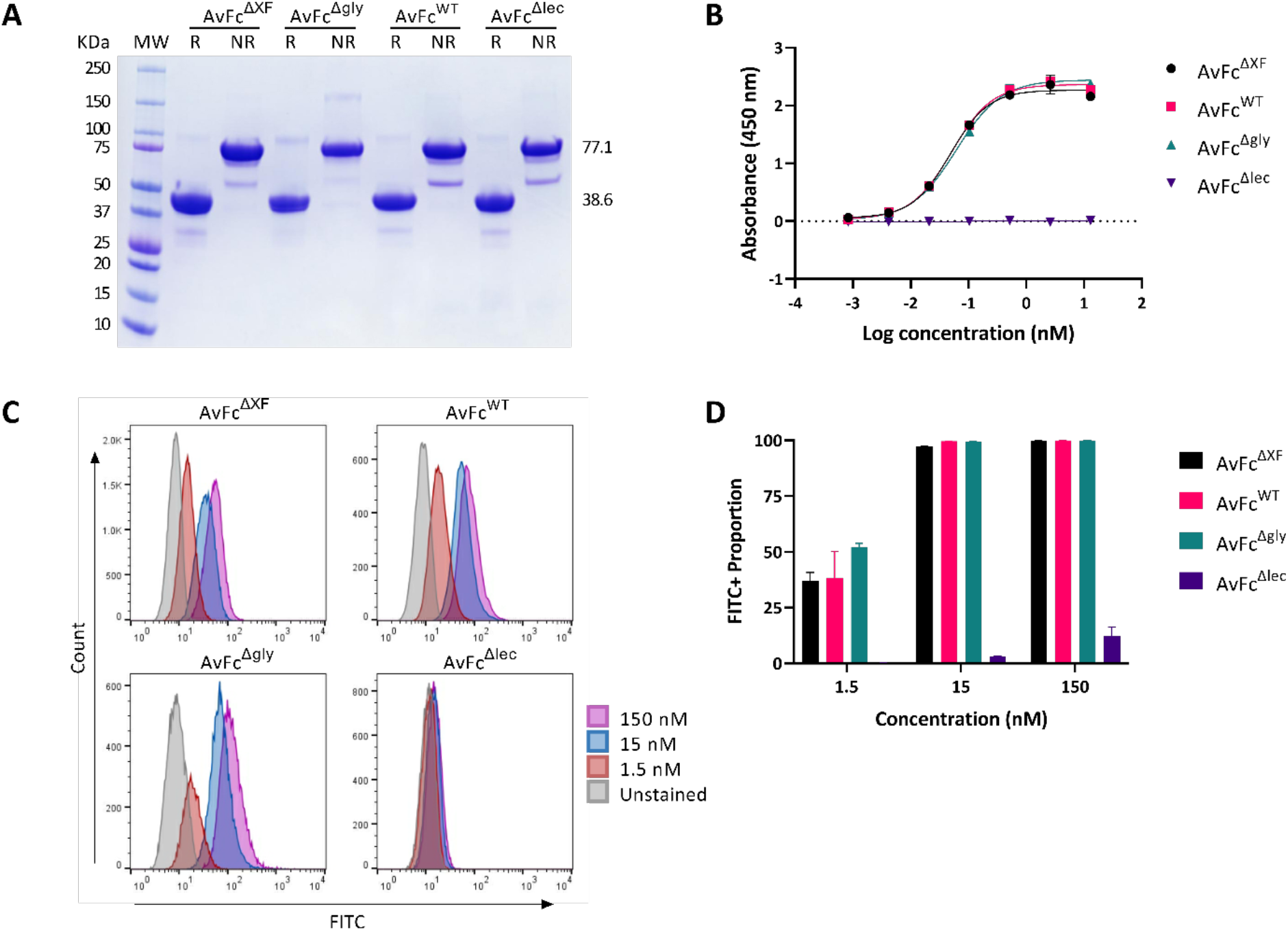
Purity and binding ability of AvFc variants do not differ significantly. (A) Purity of AvFc variants were assessed using SDS-PAGE stained with Coomassie Brilliant Blue under reducing and non-reducing conditions followed by densitometry analysis. AvFc variants appear identical according to molecular weight (≈ 38.6 kDa reduced, ≈ 77.1 kDa non-reduced) and homogenous in purity (≈ 95%), with the observation of potential Fc dimer fragments (≈ 50 kDa). (B) Comparison of AvFc variants’ binding ability as evaluated by HIV gp120-capture ELISA. Each data point represents the triplicate average ± SD. Binding curves of all AvFc variants were specified by a nonlinear regression analysis using the GraphPad Prism 9.2 software. The half maximal effective concentrations (EC50) of all AvFc variants were determined to be 0.048 nM, 0.051 nM, and 0.061 nM for AvFc^ΔXF^, AvFc^WT^, and AvFc^Δgly^, respectively. (C) Flow cytometry binding analysis of AvFc variants at 1.5 nM, 15 nM, and 150 nM to B16F10 cells. The proportion of FITC positive cells are shown at each concentration for all AvFc variants (D) with most variants saturating the cell surface at 15 nM.

### Glycoengineered AvFc retains its cancer-binding activity while improving FcγR affinity

We first set out to characterize the composition of the N-glycan on the C_H_2 domain of the human IgG1 Fc region of AvFc produced in WT or ΔXF plants (Table 1, Figure S1). By performing HPLC analysis of the C_H_2 glycan we found that AvFc expressed in WT plants contains a relatively large proportion of high mannose glycans (60.5%), with Man9 being the predominant form, followed by the expected plant-derived glycans containing both β1,2-xylose and α1,3-fucose (33.0%) and a small amount of complex glycans (6.5%). Conversely, AvFc expressed in ΔXF plants appeared to be devoid of plant-derived glycans and instead contained mostly complex glycans, primarily the GnGn glycoform (54.2%), which lacks core xylose and fucose and contains terminal N-acetylglucosamine residues, and a similarly high proportion of high-mannose glycans (40.0%).

**Table 1.** Glycan analysis of AvFc produced in WT and ΔXF *N. benthamiana* plants.

| Glycan type | Structure | AvFc<sup>WT</sup> (%) | AvFc<sup>ΔXF</sup> (%) |
| --- | --- | --- | --- |
| Plant | M3X | 1.1 | - |
|  | M3XF | 11.9 | - |
|  | GnM3XF | 3.5 | - |
|  | Gn2M3X | 2.5 | - |
|  | Gn2M3XF | 14.1 | - |
| High mannose | M7A | 2.4 | 5.3 |
|  | M7B | 5.0 | 3.3 |
|  | M8 | 18.0 | 14.8 |
|  | M9 | 35.2 | 16.6 |
| Hybrid/Complex | GnM3 | - | 1.1 |
|  | GnM4 | - | 0.3 |
|  | GnM5 | - | 3.5 |
|  | Gn2M3 (GnGn) | 6.5 | 54.2 |
| | Gal(β1,3-)Gn2M3 | - | 0.9 |
| Totals | Plant | 33.0 | - |
|  | High mannose | 60.5 | 40.0 |
|  | Complex | 6.5 | 60.0 |

Determination of the glycan composition of the single CH2 N-glycan was made using HPLC. Following expression in WT plants, AvFc displays mostly high mannose glycans followed by plant type glycans (primarily those containing both core β1,2-xylose and α1,3-fucose) and a small percentage of complex type glycans. AvFc expressed in ΔXF plants is entirely devoid of plant-specific glycans and instead consists primarily of complex glycans (primarily GnGnM3), as expected, with a significant proportion of high mannose glycans remaining (though less than when expressed in WT plants).

Next, we tested whether altering the glycosylation pattern of AvFc affected its ability to recognize cancer cells or induce Fc-mediated effector functions *in vitro.* Flow cytometry of AvFc variants binding to B16F10 cells showed that changes to the Fc glycans did not significantly impact cancer-cell binding kinetics (Figure 1C-D), with saturation of the cell surface occurring at ≈ 15 nM for AvFc^ΔXF^, AvFc^WT^, and AvFc^Δgly^. Similarly, Fc modifications did not significantly impact binding to the highly-mannosylated HIV glycoprotein gp120 as determined by ELISA (Figure 1B), with EC50 values of 0.048, 0.051, and 0.061 nM for the AvFc^ΔXF^, AvFc^WT^, and AvFc^Δgly^ variants respectively. For the AvFc^Δlec^ variant no binding to gp120 could be measured, though minor peripheral binding to B16F10 cells was observed. Despite all of the variants except AvFc^Δlec^ recognizing the cancer cell surface to similar degrees, kinetic analysis of binding to FcγRs by surface plasmon resonance (SPR) showed that the AvFc^ΔXF^ variant had ≈ 2-fold increased affinity to human FcγRI (hFcγRI), ≈ 4-fold increased affinity to human FcγRIIIa (hFcγRIIIa), and ≈ 5.5-fold increased affinity to mouse FcγRIV (mFcγRIV) compared to AvFc^WT^ (Figure 2). As expected, the aglycosylated AvFc^Δgly^ variant had little-to-no affinity observed for any of these receptors and hence a K_D_ value could not be determined. The impact of this increased affinity was assessed in an *in vitro* ADCC reporter assay, wherein activation of hFcγRIIIa on engineered Jurkat effector cells in the presence of an antibody and a target cell leads to the expression of luciferase, serving as a surrogate for Fc-mediated cell death (Oh et al., 2021). In this assay, using B16F10 as the target cell, neither AvFc^Δgly^ or AvFc^Δlec^ were capable of inducing luciferase expression, likely due to the lack of significant affinity to hFcγRIIIa or to B16F10 cells (Figure 3A). As hypothesized, incubation with AvFc^ΔXF^ resulted in the highest level of luciferase induction (≈ 5.5-fold over background, EC50 = 2.75 nM) while AvFc^WT^ showed only a moderate level of induction (≈ 2.9-fold over background, EC50 = 13.78 nM), indicating that increased hFcγRIIIa affinity has functional consequences that could potentially impact

**Figure 2.**
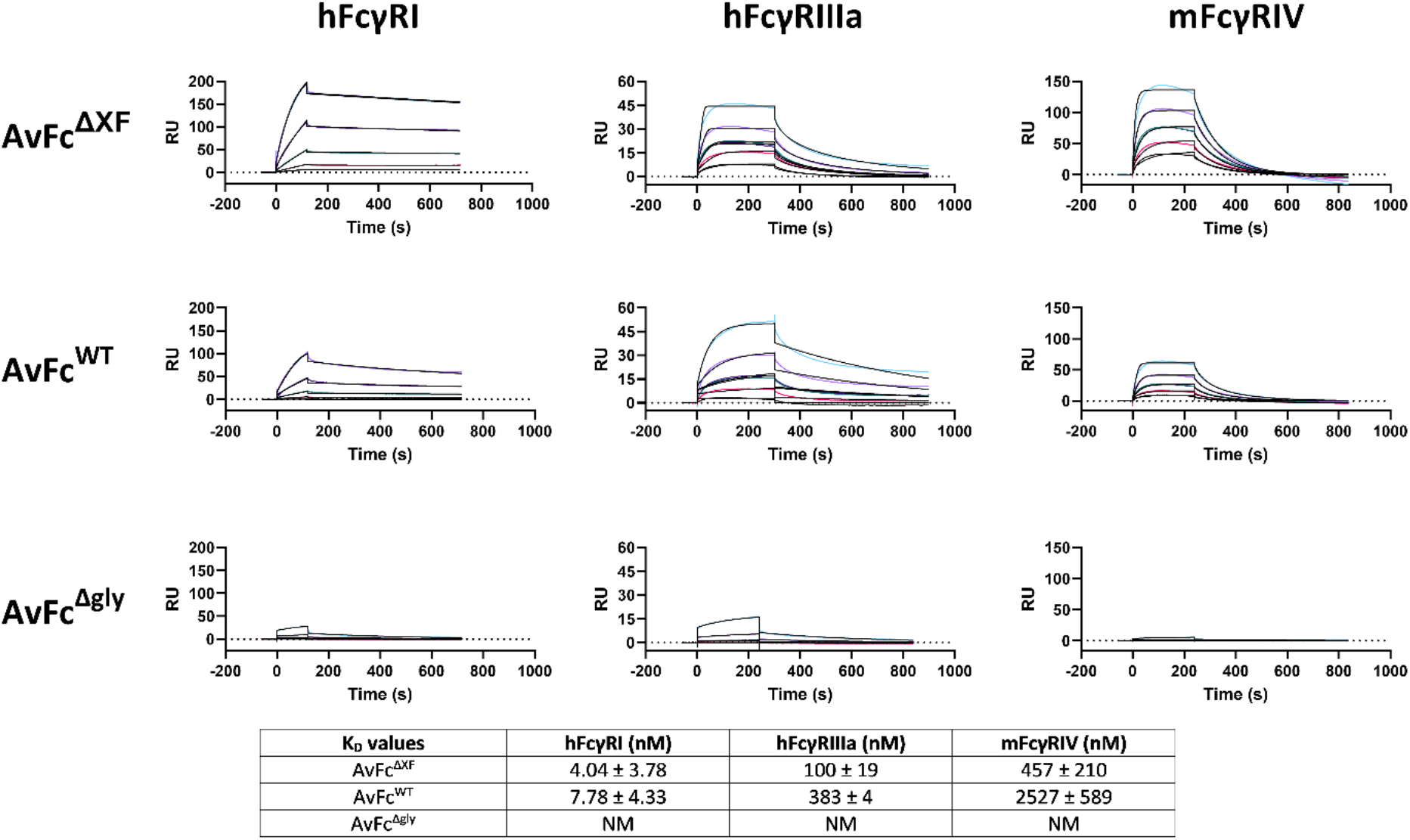
AvFc^ΔXF^ exhibits higher affinity to FcγRs from humans and mice. Shown are representative sensorgrams illustrating the association and dissociation kinetics of AvFc to the various FcγRs. Kinetics of binding to the high-affinity receptor hFcγRI are characterized by rapid association and slow dissociation, which results in low nanomolar K_D_ values. AvFc^ΔXF^ had 1.9-fold increased affinity to this receptor compared to the WT variant. For hFcγRIIIa and mFcγRIV, binding kinetics were generally characterized by rapid association and dissociation resulting in high nanomolar K_D_ values. AvFc^ΔXF^ had 3.8-fold higher affinity to hFcγRIIIa and 5.5-fold higher affinity to mFcγRIV than the WT variant. No affinity could be measured for the Δgly variant. NM = not measurable.

**Figure 3.**
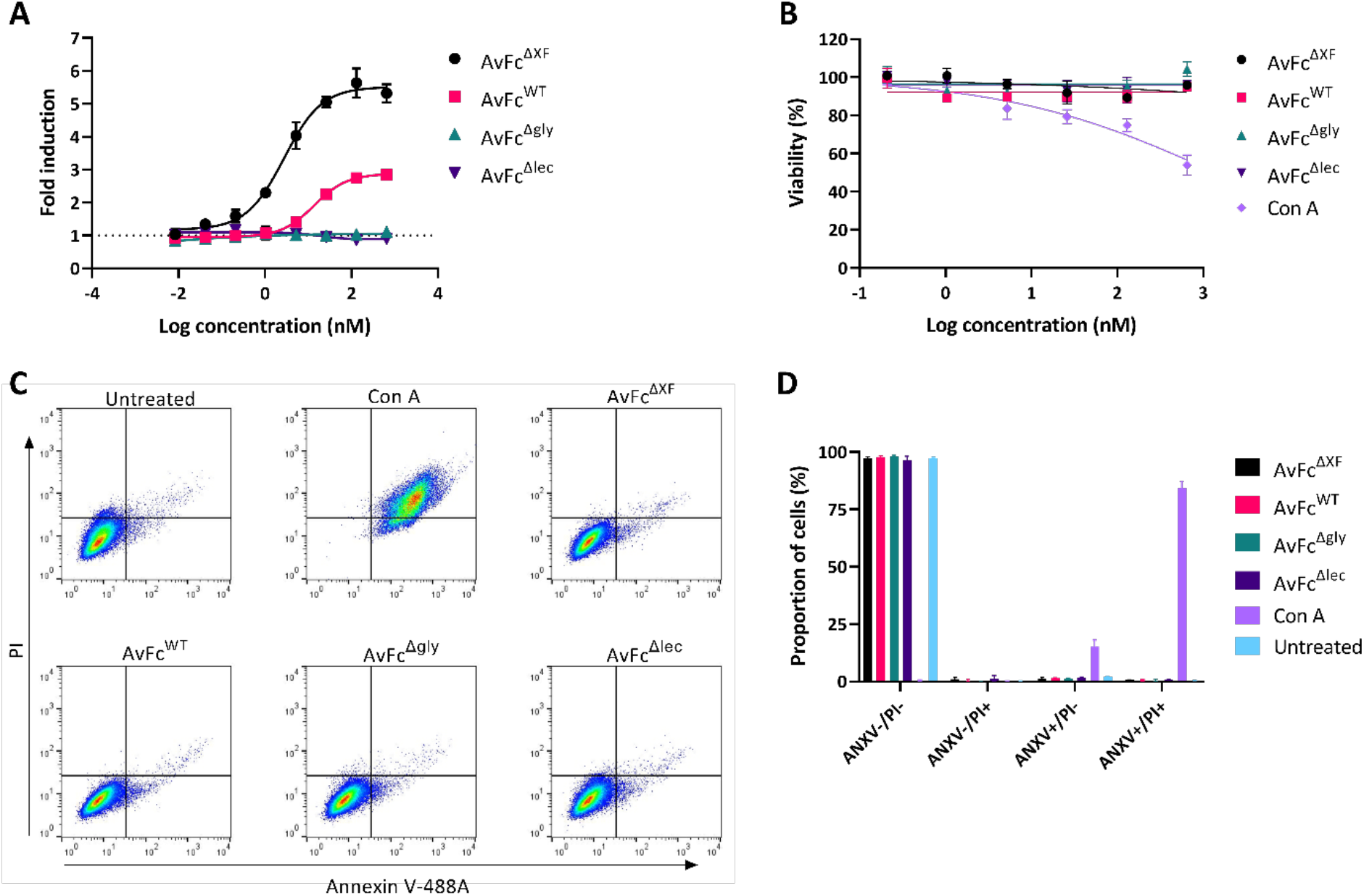
AvFc binding to B16F10 cells induces ADCC but not direct cytotoxicity. (A) The ability of AvFc variants to induce ADCC against B16F10 cells was assessed using a standardized reporter-cell-based luciferase assay. Each variant was tested in triplicates beginning at 650 nM with 1:5 serial dilutions. Each data point represents the mean ± SD while the curves were fit with a 4-parameter non-linear regression model analysis in GraphPad Prism 9.2. (B) Effects of AvFc variants and concanavalin A (conA) on B16F10 cells was measured using an MTS assay. All AvFc variants and conA were tested in triplicates beginning at 650 nM with 1:5 serial dilutions and incubated with B16F10 cells for 48 hours. Each data point represents the mean ± SD while the dose-response curves were fit using non-linear regression in GraphPad Prism 9.2. (C) Cell viability was detected using an annexin V/propidium iodide apoptosis assay using AvFc variants and conA at 650 nM. Flow cytometry dot plots with propidium iodide plotted against annexin V are shown. (D) Graphical representation of annexin V/propidium iodide assay results suggest that AvFc variants do not cause direct cytotoxicity in comparison to a known cytotoxic lectin.

AvFc’s activity *in vivo.* We further noted that AvFc, by binding to the cell surface alone, failed to induce cell death or inhibit cell proliferation after 48 hours of incubation with B16F10 cells as determined by an MTS viability assay (Figure 3B) and annexin V/propidium iodide staining (Figure 3C-D). This is in sharp contrast to concanavalin A, which is a known cytotoxic lectin that results in significant cell death when incubated with B16F10 cells (Figure 3B-D) (Kulkarni and McCulloch, 1995). Taken together, these results suggest that AvFc may exert an anti-cancer effect against B16F10 tumors primarily through Fc-mediated effector functions. To evaluate this, we opted to directly compare the AvFc^ΔXF^and AvFc^Δgly^ variants in the flank tumor model as they would represent the extreme ends of the spectrum of Fc-mediated activities of AvFc, which would effectively reveal the extent to which Fc functions are necessary for AvFc’s *in vivo* antitumor activity against B16F10 tumors.

### AvFc^ΔXF^ demonstrates activity against B16F10 flank tumors *in vivo*

To test the *in vivo* tumor-targeting capability of AvFc, we employed PET/CT imaging of live mice with established B16F10 flank tumors. Animals were injected with 1×10^6^ cells subcutaneously in the hind right flank and imaged after 10 days, at which point 3.7 MBq of a radiolabeled ^64^Cu-NOTA-AvFc was administered intravenously. Analysis of the imaging data shows that AvFc strongly accumulates inside the tumor (Figure 4D), with some additional signal observed in the liver, spleen, and bladder. These organs are the primary sites of protein metabolism, and as such background signal in these organs is commonly observed in live animal PET imaging using antibody probes (Bridgwater et al., 2020).

**Figure 4.**
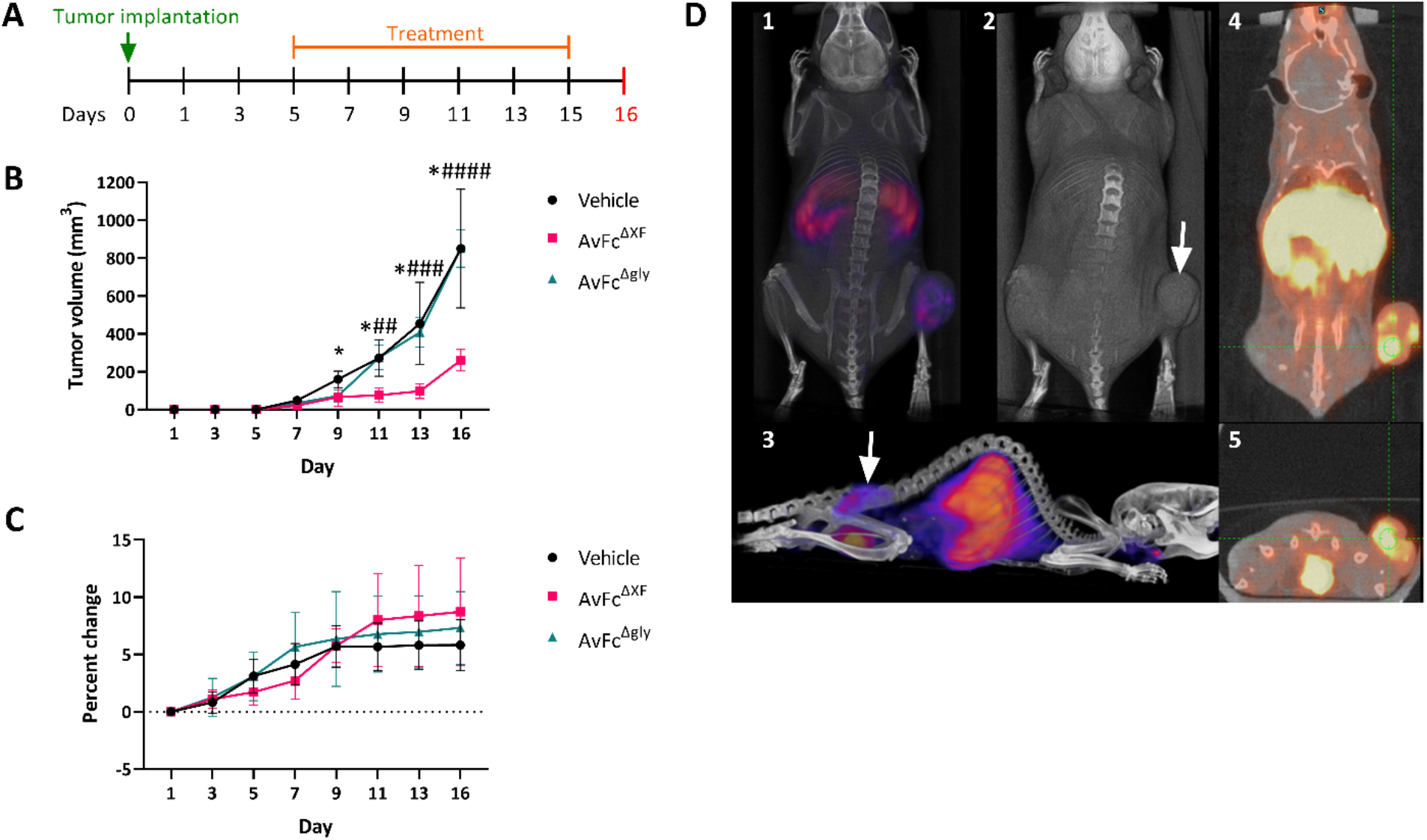
Fc effector functions are required for AvFc’s efficacy in the B16F10 flank tumor model. (A) The B16F10 flank tumor model in C57bl/6 mice (n = 5/group). On day 0, 100,000 B16F10 cells were implanted subcutaneously in the left flank. Treatment with 25 mg/kg of AvFc^ΔXF^ was initiated on day 5 and intraperitoneal dosing continued Q2D until the sixth dose on day 15. Animals were euthanized on day 16. (B) Tumor volumes were recorded every other day starting on day 1 post-implantation to day 13 with a final tumor measurement before euthanasia on day 16. Each data point represents the group mean ± SD. Tumor volumes between groups were compared with two-way ANOVA followed by Tukey multiple comparison tests (* p<0.05 between AvFc^ΔXF^ and vehicle; # p<0.05; ## p<0.01; ### p<0.001; #### p<0.0001 between AvFc^ΔXF^ and AvFc^Δgly^). (C) Comparison of body weights between groups during the study, shown as percent change from day 0 weight. No significant differences were noted between groups. (D) Representative PET/CT image of C57bl/6 mice with colocalization of radiolabeled ^64^Cu-NOTA-AvFc with B16F10 flank tumors. (D1) Whole body PET/CT, dorsal view. Signal is clearly visible within the tumor and in the liver. (D2) Whole body CT, dorsal view. Tumor is indicated by white arrowhead. (D3) Whole body PET/CT, lateral view. In addition to signal in liver and tumor, some signal is seen under the jaw, possibly corresponding to the thyroid. (D4) Coronal slide view, PET/CT scan. Tumor is indicated by the green crosshairs, which also correspond to the transverse slide view in subpanel 5. (D5) Transverse slide view, PET/CT scan.

To assess treatment with AvFc in this model, 100,000 B16F10 cells were injected subcutaneously into the hind left flank of the animal. Tumor sizes were measured every other day beginning the day after implantation (Figure 4A). Intraperitoneal treatment with 25 mg/kg of AvFc^ΔXF^, begun 5 days post tumor implantation and dosed every other day (Q2D), significantly slowed the growth of the tumors beginning from day 9 compared to both the vehicle and AvFc^Δgly^ groups. No overt toxicity was observed as determined by body weight changes over time (Figure 4B-C). AvFc^Δgly^, on the other hand, had no effect on tumor growth over time while maintaining a similar safety profile.

### AvFc’s sugar-binding activity is essential for protection against B16F10 lung metastasis

We set out to further characterize AvFc’s activity using a metastatic B16F10 melanoma challenge model, comparing AvFc^ΔXF^ with the non-sugar-binding mutant AvFc^Δlec^. In this model, 250,000 cells were injected intravenously via the tail vein followed by co-treatment with 25 mg/kg of AvFc^ΔXF^, which began on day 0 and continued Q2D for a total of 6 doses (Figure 5A). In this treatment regimen AvFc^ΔXF^ was found to significantly reduce the lung tumor burden by more than 3-fold (p=0.0009) while the non-sugar-binding mutant AvFc^Δlec^ offered no protection at the same dose (p>0.9999, Figure 5B). As described above in the flank tumor model, repeated administration of AvFc was not associated with any overt toxicity or body weight effects (Figure 5C). Based on these data, we concluded that AvFc has potent *in vivo* activity against B16F10 melanoma, and its antitumor mechanism appears to be both dependent on Fc-mediated effector functions and high-mannose binding but not by direct cytotoxicity.

**Figure 5.**
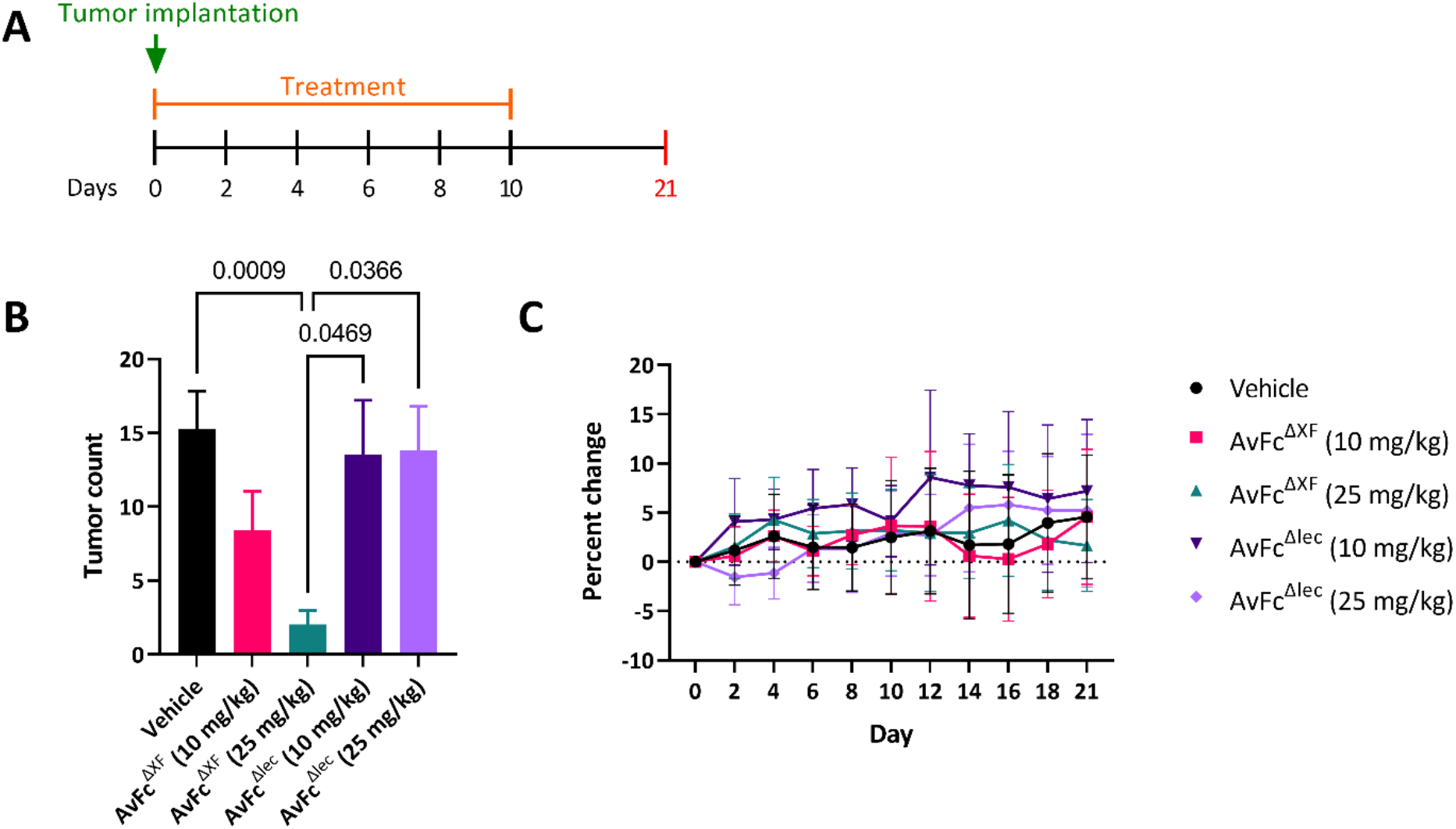
Protection against metastatic B16F10 challenge requires glycan-binding. (A) The B16F10 metastasis model in C57bl/6 mice (vehicle n=39; AvFc^ΔXF^10 mg/kg n=18; AvFc^ΔXF^25 mg/kg n=28; AvFc^Δlec^ 10 mg/kg n=10; AvFc^Δlec^ 25 mg/kg n=10) with treatment of AvFc^ΔXF^ or AvFc^Δlec^ at 10 or 25 mg/kg. On day 0, 250,000 B16F10 cells were injected intravenously followed by treatment with AvFc variants as described. Treatment continued Q2D for a total of 6 doses. Animals were euthanized on day 21. (B) Tumor count as determined by average number of visible tumor nodules on the lungs is shown for each group ± SEM. Treatment with AvFc^ΔXF^ at 25 mg/kg significantly reduced the presence of lung nodules in comparison to AvFc^Δlec^ at 10 mg/kg (p = 0.0469) and 25 mg/kg (p = 0.0366), as well as the vehicle group (p = 0.0009). Significance was determined by Kruskal-Wallis analysis followed by Dunn’s multiple comparisons in GraphPad Prism 9.2. (C) Changes in body weight over time, shown as average percent change ± SD. No significant changes in body weight were observed as determined by 2-way ANOVA.

### Pretreatment with AvFc increases survival in the B16F10 flank tumor model

As a foreign protein, AvFc administration in mice would result in the generation of anti-drug antibodies (ADAs), which in theory could compromise the safety and efficacy of the drug (Pratt, 2018). In order to address the consequence of ADA generation on the activity of AvFc *in vivo,* we modified the flank tumor model in which the experiment was divided into 2 phases: (i) a pretreatment phase, where groups of animals would receive either AvFc^ΔXF^ or a vehicle before tumor implantation to generate ADAs, and (ii) a treatment phase, where groups of animals would receive AvFc^ΔXF^ or vehicle following tumor implantation as described above. For this study we employed survival as the primary endpoint, which was defined as the time from tumor implantation until they reached a volume of 1500 mm^3^, at which point animals were euthanized and blood and other organs taken for analysis. Pretreated animals received 6 doses of AvFc^ΔXF^ at 25 mg/kg Q2D followed by an 11-day waiting period before tumor implantation (Figure 6A). To confirm the presence of ADAs, serum titers were measured by AvFc-binding ELISA at three points: just prior to tumor implantation, at the beginning of the treatment phase of the study, and following euthanasia of the animals. The results of these assays are reported in Figure 6B. Before the pretreatment, all animals had titers at or near the lowest dilution tested (1:50), possibly due to a small matrix effect caused by the presence of mouse hemoglobin on the ELISA (Reeder et al., 2004). By day 20, all of the pretreated animals had measurable ADA titers, with values between 10^4^-10^5^ that continued to increase through the end of the study at varying rates. Animals that received AvFc^ΔXF^ only during the treatment-phase of the study also generated a robust ADA response by the time of euthanasia, between 10^4^ and 10^6^. By far, the pretreated and AvFc-treated group generated the largest ADA response, albeit a more variable one, with titers between 10^4^ and 10^8^ at euthanasia.

**Figure 6.**
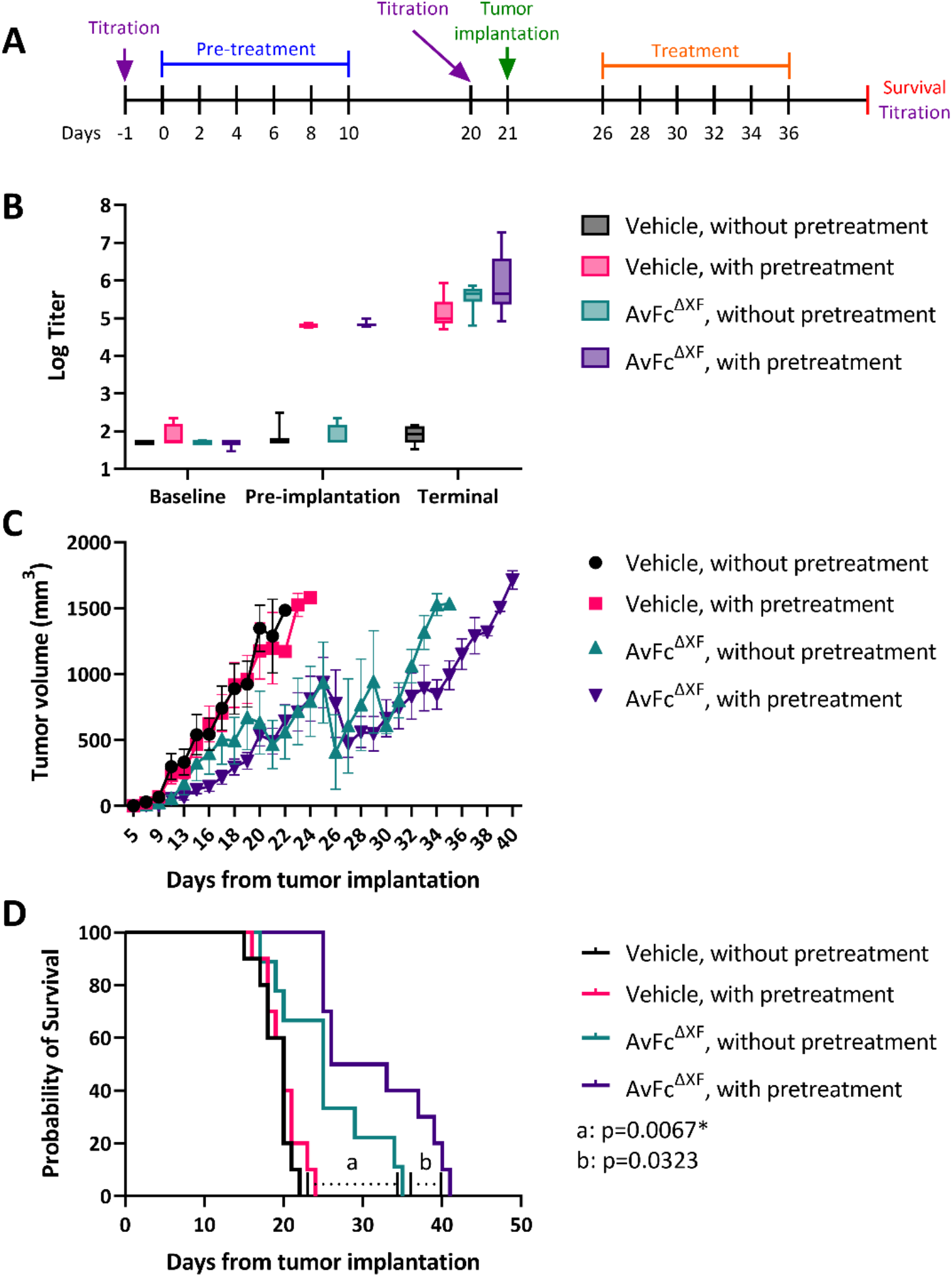
Impact of pretreatment on the activity of AvFc *in vivo.* (A) The B16F10 flank tumor model with pretreatment of AvFc^ΔXF^ in C57bl/6 mice (n = 10/group). Pretreatment groups received 6 intraperitoneal doses of AvFc^ΔXF^ at 25 mg/kg Q2D prior to flank tumor challenge with 100,000 B16F10 cells on day 21. Beginning on day 26, treatment groups received 6 doses of AvFc^ΔXF^using the same dosing parameters as described for pretreatment. Animals were euthanized at a tumor volume of 1500 mm^3^. (B) Baseline, pre-implantation, and terminal anti-drug antibody (ADA) titers for all experimental groups. The minimum, median, and maximum titer values are indicated. Baseline titers were likely affected by a matrix effect from the mouse serum. Day 20 pre-implantation titers of pretreatment animals resulted in values between 10^4^-10^5^ that increased through the terminal titers at differing rates. ADA titers of animals that received only treatment post-implantation revealed values between 10^4^ and 10^6^ at the time of the terminal titers. (C) Flank tumor volumes were recorded with euthanasia taking place at 1500 mm^3^. Each data point represents the mean tumor volume ± SEM. (D) Survival rates for all experimental groups. Statistical significance was determined using the Mantel-Cox log-rank test (p<0.01). Significance was observed between non-pretreated, AvFc^ΔXF^ dosed group survival and both vehicle-treated groups (p=0.0067 vs. non-pretreated animals, p=0.0083 vs. pretreated animals). Additional significant relations were identified between pretreated, AvFc^ΔXF^ dosed group survival and both vehicle-treated groups (p<0.0001 vs. each vehicle-treated group).

The effects of pretreatment on animal survival in this model are summarized in Figure 6C-D. Pretreatment with AvFc^ΔXF^ had no effect on the survival of vehicle-treated animals compared to non-pretreated animals (p=0.3049). As was previously observed, treatment with AvFc^ΔXF^ of non-pretreated animals resulted in delayed tumor growth and increased survival compared to vehicle-treated animals (p=0.0067 vs. non-pretreated animals, p=0.0082 vs. pretreated animals). Interestingly, compared to the non-pretreated AvFc^ΔXF^-treated group, pretreatment with AvFc^ΔXF^ extended the median survival time by nearly 5 days (25 vs. 29.5), though this effect failed to reach statistical significance after correction for multiple comparisons (p=0.0323). These data reveal that ADAs generated against AvFc do not appear to neutralize the drug and make it ineffective, nor did they present any obvious safety concerns over the course of the study with no major adverse events or body weight effects noted. On the contrary, it seems that the presence of ADAs may have increased the anti-tumor activity of AvFc^ΔXF^.

## DISCUSSION

Plant-based recombinant expression systems have found some success as rapid, robust, and scalable alternative manufacturing platforms for pharmaceutical proteins (Arntzen, 2015; Daniell et al., 2015; Politch et al., 2021; Ward et al., 2021; Ward et al., 2020). A useful characteristic of many of the plants used for pharmaceutical production, in particular *N. benthamiana,* is that they are generally readily amenable to engineering using modern techniques including RNA interference (RNAi), transcription activator-like effector nucleases (TALENs), zinc-finger nucleases, and CRISPR/Cas9 (Casacuberta et al., 2015; Feng et al., 2013; Khan et al., 2017; Petolino, 2015). Therefore, *N. benthamiana* can be exploited to generate glycoengineered variants of biologics that have higher levels of activity *in vivo* and may obviate some safety concerns regarding the presence of plant N-glycans. Removal of either the human α1,6- or plant α1,3-linked core fucose residues through genetic engineering of the expression host has long been known to dramatically increase the affinity of monoclonal antibodies for FcγRs, especially hFcγRIIIa, and improve their activity both *in vitro* and *in vivo* (Nimmerjahn and Ravetch, 2005; Shields et al., 2002). Thus, for therapeutic antibodies or other Fc-bearing molecules such as AvFc where ADCC is a major mechanism of action, such a modification would be highly valuable. In this study, we show that expression of our candidate anti-cancer immunotherapeutic AvFc in glycoengineered ΔXF plants results in the total loss of plant-derived glycans, with the predominant glycoform being the truncated, “humanized”, GnGn form (Table 1, Figure S1). However, compared to mAbs expressed in WT or ΔXF plants, AvFc displays some idiosyncrasies with regards to its glycan composition.

The first of these is that AvFc^WT^ contains few plant-derived glycans, with glycoforms containing β1,2-xylose and α1,3-fucose (XF) representing only 33% of the total glycan population compared to upwards of 90% for most plant-expressed mAbs (Table 1) (Bardor et al., 2003; He et al., 2014; Lee et al., 2013; Schähs et al., 2007). The second is that both AvFc^WT^ and AvFc^ΔXF^ display relatively large proportions of high-mannose glycans, which are typically only present on mAbs in very small amounts when expressed in plants (Bardor et al., 2003; He et al., 2014; Lee et al., 2013; Schähs et al., 2007). These two observations may indeed be somewhat linked, as an overabundance of high-mannose glycans on recombinant proteins can be the result of extended residency or accumulation in the endoplasmic reticulum, such as is seen when antibodies are tagged with the ER-retention signal KDEL (Triguero et al., 2005). Given that AvFc is a high-mannose-binding lectibody, it is highly possible that it forms complexes with itself or with other ER-resident glycoproteins during expression, preventing export to the Golgi apparatus for further processing and resulting in an accumulation of high-mannose glycans. Similarly high levels of high-mannose glycans have been identified on mAbs with atypical structures such as mono- and multivalent single-chain variable fragments, which are hypothesized to be retained in the ER due to prolonged interaction with BiP in the absence of the light chain constant region (Feige et al., 2009; He et al., 2014). However, similar findings have not been reproduced for Fc fusion proteins produced in mammalian cells or in plants (Bongers et al., 2011; Jones et al., 2007; Keck et al., 2008; Lu et al., 2012; Lynaugh et al., 2013; Nagels et al., 2012; Schriebl et al., 2006). It should be noted that AvFc manufactured in CHO cells also has increased levels of high-mannose glycans compared to normal mAbs, though there are fewer and the predominant form is Man5 (data not shown), indicating a greater degree of processing but also suggesting that the increase is due to a property of AvFc and not the production host. The increased presence of high-mannose glycans may also help to partially explain the relatively low half-life of AvFc^ΔXF^ in mice and rhesus macaques (≈ 22 and ≈ 28 hrs, respectively, as reported in (Dent et al., 2021; Hamorsky et al., 2019)) compared to normal mAbs, as antibodies and other Fc-bearing molecules containing high-mannose glycans are more rapidly cleared from the organism by the immune system (Goetze et al., 2011). AvFc^ΔXF^ also had a longer half-life in female mice than AvFc^WT^ (≈ 14 hours for the WT variant, data not shown vs. ≈ 18.5 hrs for the ΔXF variant, Figure S2 and (Dent et al., 2021)), possibly due to the reduction in high-mannose glycan content compared to the WT (Table 1). Additionally, we have previously shown that AvFc has lower affinity to FcRn compared to normal human IgG1, which also impacts half-life (Hamorsky et al., 2019). Improvements to half-life could be made through the introduction of amino acid substitutions that improve FcRn affinity such as those described by Mackness et al. (Mackness et al., 2019). Work is being done to elucidate the cause of the increase in high-mannose, including by assessing the glycan profile of the non-sugar-binding mutant AvFc^Δlec^, which may not interact with itself or other proteins in the ER.

It is well known that removal of the core fucose residues increases the ADCC activity of mAbs (Liu et al., 2015). Our results show that AvFc^ΔXF^, which lacks both core xylose and core fucose, has similarly higher affinity to both hFcγRIIIa (and its murine equivalent mFcγRIV) and hFcγRI in addition as determined by SPR (Figure 2). Functionally, removal of plant-derived β1,2-xylose and α1,3-fucose made the AvFc^ΔXF^ variant nearly twice as efficacious and 5 times more potent in the ADCC reporter assay against B16F10 cells, while neither the Δlec nor Δgly AvFc variants were capable of inducing ADCC (Figure 3A).

Results from the *in vitro* assays (Figure 3) and the B16F10 challenge model (Figure 4) suggest that the antitumor activity of AvFc is almost exclusively attributable to the induction of Fc-mediated effector functions, in particular ADCC. As shown in Figure 3, AvFc induced high levels of ADCC without directly causing cell death as determined by both an MTS assay and annexin V/propidium iodide staining, with the AvFc^ΔXF^ variant producing a much higher ADCC response than AvFc^WT^. In addition, we did not find that AvFc inhibited B16F10 migration (data not shown), demonstrating that binding alone is likely not sufficient for AvFc to exert activity against B16F10 cells. However, we have also recently reported that AvFc can inhibit the migration of H460 and A549 human lung cancer cells in addition to inducing ADCC against these cell lines (Oh et al., 2021). The discrepancy between the findings in the present study with B16F10 cells and the previous one with H460 and A549 may be partly explained by specific cell-surface glycoproteins targeted by AvFc; for example, in the previous study we showed that the lectibody’s binding to EGFR and IGF1R led to the inhibition of receptor phosphorylation and downstream signaling (Oh et al., 2021). Thus, we speculate that the collective antitumor mechanisms of AvFc may be dependent on the characteristics of cancer cells targeted although Fc-mediated activity is likely the lectibody’s primary mode of action. We also reported in this study that administration of AvFc^ΔXF^ but not AvFc^Δgly^ delayed the growth of B16F10 flank tumors *in vivo* (Figure 4). AvFc^ΔXF^ was also found to significantly reduce the number of tumor nodules in mouse lungs in the B16F10 metastasis model while the non-sugar-binding mutant AvFc^Δlec^ did not (Figure 5). Combined, these results suggest that, at least in the B16F10 model, both sugar-binding activity and Fc-mediated effector functions are necessary for AvFc’s anti-cancer activity, and direct cell-killing likely does not occur.

A potential concern when administering foreign proteins as therapeutics is immunogenicity, which has the potential to limit drug efficacy after repeated dosing and can result in serious adverse events related to hypersensitivities, owing to the induction of ADAs and immunological memory to the drug (Chirmule et al., 2012). One of the more striking findings in this study was that pretreatment with AvFc and generation of ADAs did not negate AvFc’s activity (Figure 6). On the contrary, it appears that the presence of ADAs may have improved its activity, resulting in an increase in median survival time (defined as the time from tumor implantation to the time the tumor reached a volume of 1500 mm^3^) from 25 to 29.5 days compared to the AvFc^ΔXF^-treated group without preexisting ADAs. While this increase was not statistically significant after correcting for multiple comparisons (p=0.0323), the data show a clear trend that at the very least indicates that the ADA response to AvFc did not undermine the drug efficacy in this model. At most, it demonstrates that the presence of ADAs may actually have some benefit, though the reason for this observation remains a subject of speculation at this point.

In recent years, it has become increasingly evident that tumors are highly adept at managing the local immune microenvironment, converting it from an immunogenic to an immunosuppressive environment (Binnewies et al., 2018; Hinshaw and Shevde, 2019; Ning et al., 2021). This conversion makes tumors more aggressive and allows them to better invade the surrounding tissues and metastasize, leading to poorer clinical outcomes (Saleh and Elkord, 2020). This fact has led to the institution of a novel paradigm of cancer immunotherapy whose objective is to target not the tumor but the host immune response, helping to convert immunologically “cold” tumors to immunologically “hot” ones that can be better treated (Pitt et al., 2016). The most prominent examples of this are the checkpoint inhibitors that target the inhibitory receptors PD-1 and CTLA-4 on T cells, preventing the tumor-initiated deactivation of cytotoxic T lymphocytes in the local microenvironment leading to a better anti-tumor immune response and better treatment outcomes (Abdou et al., 2020). Other approaches, such as vaccination with TAAs or administration of immunocytokines, work by increasing the anti-tumor antibody response as well as by stimulating the tumor immune microenvironment to become more inflammatory (Murer and Neri, 2019; Vermaelen, 2019).

We initially hypothesized that AvFc, as a foreign protein that selectively recognizes tumor cells, may work in a similar manner by stimulating the local immune response to the tumor and increasing tumor antigen presentation and generation of anti-tumor antibodies (ATAs). A preliminary flow cytometry experiment using B16F10 cells stained with serum from pretreated and non-pretreated animals from this study and a goat anti-mouse IgG-FITC conjugate demonstrated a slight but non-significant increase in anti-tumor antibodies as inferred from the increased number of cells with a mean fluorescence greater than the background. (Figure S3D). We also performed a modified ADCC assay by spiking AvFc^ΔXF^ into solutions of pooled serum obtained from animals during the immunogenicity study shown in Figure 6 to assess whether or not ADAs or ATAs impact ADCC induction in the reporter cell assay (Figure S3A-C). With this method we found that while serum taken from any time point did not induce any luciferase expression alone (Figure S3A-B), the addition of AvFc^ΔXF^ to the serum resulted in an increase in activity significantly greater than is induced by AvFc^ΔXF^ alone (Figure S3C). Furthermore, the effect on induction is greatest with AvFc^ΔXF^ spiked into serum from animals that received both pretreatment and treatment with AvFc^ΔXF^ (Figure S3C). Taken together, these data suggest that perhaps the increased activity resulting from pretreatment is not necessarily due to an increase in anti-tumor antibodies but by both an additive or synergistic effect on ADCC between ADAs, ATAs, and AvFc in addition to changes in the cellular composition of the immune microenvironment following treatment. Such a phenomenon has been observed following the successful treatment of murine ID8 ovarian cancer by cowpea mosaic virus-like particles, which was similarly not affected by the generation of ADAs and resulted in substantial changes to the immune cell composition towards a more inflammatory phenotype (Shukla et al., 2020; Wang et al., 2019a). To explore this hypothesis, we have performed a preliminary flow cytometry analysis of infiltrating immune cells isolated from B16F10 tumors and have observed a significant increase in the infiltration of non-classical monocytes (Figure S4, Figure S5). Non-classical monocytes are a functionally distinct subset of steady-state monocytes that express pro-inflammatory cytokines and FcγRs, allowing them to both recruit immune cells to sites of injury or cancer and undergo antibody-dependent phagocytosis (Bharat et al., 2017; Mukherjee et al., 2015). While much of the biology of these cells remains unknown, recent studies have shown that they are important for the control of metastasis by eliminating tumor cells in the vascular beds (Hanna et al., 2015; Thomas et al., 2016). While these results are preliminary, they corroborate our assertion that Fc-mediated functions are crucial to the anti-cancer activity of AvFc and suggest that the recruitment of FcγR-bearing cells into the tumor microenvironment may play a significant role in its mechanism of action.

In conclusion, the present study has demonstrated the successful glycoengineering and activity of a novel immunotherapeutic drug, AvFc, which is a lectibody targeting cancer-associated high-mannose glycans. This glycoengineered AvFc, which lacks plant-derived glycans (in particular the core α1,3-fucose), induces more a potent ADCC response *in vitro* and delays the growth of murine B16F10 melanoma in both a flank tumor model as well as a model of metastasis *in vivo.* Additionally, pretreatment with AvFc and generation of ADAs did not negate AvFc’s activity and indeed may have increased it through a yet undetermined mechanism. These findings further substantiate the notion that high-mannose glycans may be a useful druggable biomarker in cancer therapy, and that glycoengineering is a powerful strategy to improve the antitumor activity of AvFc.

## EXPERIMENTAL PROCEDURES

### Animal Care

Animal studies were conducted with the approval of the Institutional Animal Care and Use Committee of the University of Louisville. All animals were housed in small groups in a facility with a 12 hr day/night cycle and given access to standard food and water *ad libitum.* Animals were acclimated for at least one week following their arrival to the facility before the beginning of each study.

### Cell Culture

B16F10 murine melanoma cells were acquired from the American Type Culture Collection (CRL-6475™, Manassas, VA) and cultured in high-glucose Dulbecco’s Modified Eagle Medium (Corning 10-013-CV, Corning, NY) containing L-glutamine and sodium pyruvate, supplemented with 10% fetal bovine serum (VWR 89510-196, Radnor, PA) and 1X penicillin/streptomycin (VWR 97063-708). Modified Jurkat ADCC effector cells expressing hFcγRIIIa and firefly luciferase downstream of an NFAT response element were obtained from Promega (G7102, Madison, WI) and cultured according to the kit protocol. All cells were incubated at 37°C with 5% CO_2_.

### Expression, purification, and endotoxin removal of AvFc and its variants

AvFc^WT^ and its variants (AvFc^Δlec^, AvFc^Δgly^, AvFc^ΔXF^) were recombinantly expressed in wild type *Nicotiana benthamiana* plants using the deconstructed tobacco-mosaic-virus-derived three-component vector system magnICON^®^ and purified using Protein A and CHT FPLC as described previously (Hamorsky et al., 2019). The lectin-deficient variant (AvFc^Δlec^) was made by introducing three point mutations to the Avaren lectin (Y32A, Y70A, Y108A), and was confirmed devoid of sugar-binding activity by gp120-binding ELISA. The aglycosylated variant (AvFc^Δgly^) was made by introducing a single point mutation (N200Q) to the lone N-glycan site on the C_H_2 domain of the human Fc region. The AvFc^ΔXF^ glycovariant was expressed in glycoengineered *N. benthamiana* (ΔXF) plants and purified in the same manner. Endotoxin was removed from purified lots using Triton X-114 phase separation after CHT chromatography and prior to final formulation and sterilization (Teodorowicz et al., 2017). Briefly, Triton X-114 was added to protein samples to a final concentration of 1% and incubated on ice for 30 minutes before heating to 37°C for 10 minutes and spinning at maximum speed for 30 minutes at 37°C. The endotoxin-free protein layer above was separated from the endotoxin-rich layer at the bottom by careful pipetting and residual detergent was removed by diafiltration. Additional Triton X-114 wash cycles were performed with 0.1% detergent followed by heating and centrifugation as before. Endotoxin levels were measured using the Endosafe PTS™ system (Charles River, Wilmington, MA). Pure endotoxin-free protein (< 0.2 EU/mg) was diafiltrated with AvFc formulation buffer (30 mM histidine pH 7.4, 100 mM sucrose, 100 mM NaCl) using centrifuge filters with a MWCO of 10,000 kDa (Millipore Sigma 2019-10, Burlington, MA), sterilized using 0.2 μm vacuum filters, and stored at −80°C. Purity of AvFc was assessed using SDS-PAGE with Coomassie Brilliant Blue staining. Gels were imaged using an Amersham Imager 600 and densitometry was performed using the GelAnalyzer software (GelAnalyzer 19.1, www.gelanalyzer.com, by Istvan Lazar, Jr., and Istvan Lazar, Sr.). Final yield was calculated in mg of purified protein per kilogram of plant tissue harvested.

### ELISA to assess gp120 binding

Recombinant envelope glycoprotein gp120 from HIV-1 (CM235, NIH ARP 12816) was coated overnight at 4°C at 0.3 μg/mL in carbonate buffer, pH 9.5. After coating, wells were blocked with PBST containing 5% non-fat dry milk for 1 hour at 37°C. AvFc variants were then incubated on the plate beginning at 13 nM with 1:5 serial dilutions for 1 hour at 37°C, followed by detection with a 1:10,000 dilution of goat anti-human IgG-HRP (Southern Biotech 2040-05, Birmingham, AL). Plates were developed for 5 minutes with TMB substrate (VWR 95059-286), with development stopped with an equal volume of 2 N sulfuric acid and plates read at 450 nm. Dose-response curves were fit with 4-parameter non-linear regression in GraphPad Prism which were used to calculate EC_50_ values.

### Glycan analysis

The N-linked glycans were released from 1 mg of purified recombinant AvFc by hydrazinolysis (Fujiyama et al., 2006). After N-acetylation with saturated sodium bicarbonate and acetic anhydride, the hydrazinolysate was desalted using Dowex 50 × 2 (Muromachi Kagaku Kogyo Kaisya, Tokyo, Japan), and lyophilized. The oligosaccharides obtained were pyridylaminated (PA) as described previously (Fujiyama et al., 2006; Kondo et al., 1990). PA-sugar chains were purified by HPLC and monitored on the basis of the fluorescence intensity (λexc = 310 nm, λem = 380). For RP-HPLC, PA-sugar chains were eluted from a Cosmosil 5C18-AR-II column (Nacalai Tesque, Kyoto, Japan) by linearly increasing the acetonitrile concentration in 0.02% trifluoroacetic acid (TFA) from 0% to 4% for 35 min at a flow rate of 0.7 mL/min. To further determine PA-sugar chain structures, LC-MS/MS analysis was performed using HPLC equipped with ion trap mass spectrometry (HCT plus, Bruker Daltonics, Bremen, Germany) as described previously (Kajiura et al., 2015).

### Flow cytometry

B16F10 cells were stained with AvFc variants at 150, 15, and 1.5 nM followed by detection with a goat anti-human Fc FITC secondary antibody (Abcam ab97224, Cambridge, UK) at 1:200 and fixation in 4% formalin. Unstained cells incubated with the secondary antibody only and AvFc^Δlec^ (150 nM) were used as controls to determine background fluorescence. To detect anti-tumor antibodies, B16F10 cells were stained with a 1:10 dilution of individual animal serum followed by detection with a 1:1000 dilution of a goat anti-mouse Fc FITC secondary (Abcam ab98716) and fixation with 4% formalin. Flow cytometry was performed on a BD FACSCalibur and all data were analyzed in FlowJo. Statistical comparisons were made with One-way ANOVA followed by multiple comparisons with Tukey’s multiple comparison test.

### ADCC Reporter Assay

B16F10 cells were plated at 10,000 cells/well on a solid white 96-well plate and incubated overnight at 37°C to allow attachment. The following day AvFc^WT^ or its variants were serially diluted (from 650 nM to 8.32 pM in 1:5 steps) in ADCC assay buffer, which consisted of RPMI-1640 medium supplemented with 1% Ultra Low IgG Fetal Bovine Serum (VWR 4100-050) and added to the cell-containing wells. Jurkat effector cells, which were also suspended in ADCC assay buffer, were then added to each well to give a total effector:target cell ratio of 15:1 (150,000 cells) and incubated overnight. In cases where the drug and effector cells were co-incubated with animal serum, assay buffer was used to dilute the serum to the indicated percentage. After incubation, the culture medium was carefully removed, and luminescence was measured on a BioTek plate reader using the Britelite Plus Reporter Gene Assay System (Perkin Elmer 6066761, Waltham, MA). Each assay included triplicates of the treatment conditions as well as a no drug control and a B16F10 cell only control. Fold luminescence induction was plotted against the log drug concentration and was calculated as the ratio between the relative luminescence units (RLUs) of the wells containing drug and the average RLU values for the no drug control. The resulting dose-response curves were fit with a 4-parameter non-linear regression model in GraphPad Prism to calculate the EC_50_.

### MTS Assay

B16F10 cells were plated at 5,000 cells per well and incubated with AvFc (beginning at 650 nM) for 48 hours. The toxic mannose-binding lectin concanavalin A was used in equimolar concentrations as a positive control. After incubation, 20 μL of MTS reagent (Abcam ab197010, Cambridge, UK) was added to each well and incubated for 4 hours, at which point the reaction was stopped by adding 10 μL of 10% SDS. Plates were then read in a BioTek plate reader at 490 nm. Percent viability was calculated relative to untreated controls and plotted against concentration. Dose-response curves were fit using non-linear regression in GraphPad Prism to calculate IC_50_ values.

### Annexin V/propidium iodide staining of apoptotic cells

B16F10 cells (1.5×10^5^ cells/well) were seeded into 6 well plates with AvFc variants or concanavalin A at 650 nM and incubated for 48 hours at 37°C. Following incubation, cells were harvested and stained with an annexin V-488A (ANXV) conjugate and propidium iodide (PI) (Thermo Fisher Science) according to Rieger et al. (Rieger et al., 2011). Briefly, cells were stained with 2.5 μg/mL annexin V conjugate and 2 μg/mL PI for 15 minutes each at room temperature in the dark. Following fixation with a 1% formalin solution, cells were incubated with 50 μg/mL RNase A and measured on a BD FACSCalibur flow cytometer. Apoptotic cells, in the early and late stages, were defined as ANXV+/PI- and ANXV+/PI+, respectively, with unstained cells used to define the quadrant gates. Data were processed and analyzed using FlowJo, and statistical comparisons between groups were made with 2-way ANOVA with p=0.05 as the threshold of significance.

### Surface plasmon resonance for FcγR binding

Surface plasmon resonance experiments were performed on a Biacore T200. For hFcγRIIIa and mouse FcγRIV (mFcγRIV) we opted for a 6xHis-capture approach. To achieve this, an anti-6xHis-tag monoclonal antibody (Thermo Fisher Scientific) at 50 μg/mL was conjugated via amine linkage to two parallel flow cells on the surface of a CM5 chip at ≈ 10,000 response units (RUs). Recombinant Fc receptors obtained from R&D Systems were captured by flowing them over the chip surface at 5 μg/mL for 60 seconds with a flow rate of 10 μL/min. A second flow cell was used for reference subtraction and was not used to capture the receptors. AvFc variants (ΔXF, WT, Δgly) were flowed over both cells at multiple concentrations with a flow rate of 30 μL/min, using an association time of 240 seconds and a dissociation time of 600 seconds. For both hFcγRIIIa and mFcγRIV, 5 concentrations of AvFc were used to measure affinity starting at 2 μM with 1:2 serial dilutions, repeating the middle concentration and including a blank cycle. Regeneration was performed by washing the chip surface for 60 seconds at a flow rate of 30 μL/min with glycine at pH 1.5. Sensorgrams were fit with a 1:1 binding model with Rmax set to fit local using the Biacore Evaluation software.

For hFcγRI, recombinant receptor was captured on the surface of an NTA-conjugated chip following a 60 second injection of Ni^2+^ at 0.5 mM. The receptor was captured by flowing at 10 μL/min at a concentration of 1 μg/mL for 100 seconds to achieve a capture level of ≈ 200 RUs. A second flow cell was left blank for reference subtraction. 5 concentrations of AvFc were used to measure affinity starting at 324 nM with 1:3 serial dilutions, repeating the middle concentration and including a blank cycle. Regeneration was performed by washing the chip surface for 60 seconds at a flow rate of 30 μL/min with 350 mM EDTA. Sensorgrams were fit with a 1:1 binding model with Rmax set to fit local using the Biacore Evaluation software.

### PET/CT imaging

The in vivo tumor-targeting property of AvFc was determined with radiolabeled AvFc in B16F10 melanoma-bearing C57bl/6 mice using small animal PET/CT. The mice (n = 2) were each subcutaneously inoculated with 1×10^6^ B16F10 cells on the right flank to generate tumors. The animals were submitted to imaging when tumor weights reached approximately 0.2 g at 10 days post-cell inoculation. Approximately 3.7 MBq of purified ^64^Cu-NOTA-AvFc was injected into each mouse via the tail vein. The mice were scanned with small animal PET and CT at 24 h post-injection. A ten-minute CT scan (MicroCAT II) was immediately followed by 30 min PET imaging on MicroPET (Siemens R4) using the same animal bed. The PET and CT data obtained were reconstructed and merged by the Siemens IRW software.

### Pharmacokinetics of AvFc^WT^ in C57bl/6 mice

A pharmacokinetic profile for AvFc^ΔXF^ was generated following a single 25 mg/kg intraperitoneal injection in female C57bl/6 mice (The Jackson Laboratory, Bar Harbor, ME). Blood was sampled at 0.5, 1, 2, 4, 8, 12, 24, and 48 hours after injection by either submandibular vein or cardiac puncture, and 4 animals per time point were used. Pharmacokinetic parameters were calculated using Certara Phoenix WinNonlin software with a non-compartmental analysis.

### B16F10 flank tumor challenge model

On day 0, 100,000 B16F10 cells in 100 μL of DPBS was injected subcutaneously into the hind left flank of each C57BL/6 mouse (n = 5/group). Intraperitoneal administration of 200 μL of AvFc^ΔXF^, AvFc^N200Q^, or vehicle (AvFc formulation buffer, see above) at the indicated dose level began on day 5 and continued Q2D until day 16. Tumor measurements were taken every other day from day 1 using digital calipers, and tumor volume was estimated as: *tumor width x tumor height^2^*.

To determine the impact of anti-drug antibodies on the efficacy of AvFc^ΔXF^ in this model, groups of animals (n = 5/group) were pre-pretreated with 6 doses of AvFc^ΔXF^ at 25 mg/kg Q2D followed by tumor implantation 11 days after the final dose (day 21). Treatment with AvFc was then performed as before, beginning on day 5 and continuing Q2D for a maximum of 6 doses. The primary endpoint was survival, defined as the time until animals reached a tumor volume of 1500 mm^3^, at which point the animal was euthanized. Blood was collected to determine anti-drug antibody titers on day −1 and day 20 via submandibular vein before pretreatment and before B16F10 implantation, and again via cardiac puncture at the time of euthanasia. Survival curves were compared using the Mantel-Cox test in GraphPad Prism. Multiple comparisons of individual survival curves were made using the Mantel-Cox test, and the significance threshold was corrected using the Bonferroni method (corrected p value threshold was 0.0083).

### Calculation of anti-drug antibody titers

Anti-drug antibody titers were measured by ELISA. AvFc^ΔXF^ was coated on a 96-well plate at 1 μg/mL overnight at 4°C, followed by blocking for 1 hour with 3% BSA-PBST at room temperature. Mouse serum was then plated at an initial minimum dilution of 1:50 and serially diluted further with 1:10 dilutions, followed by a 2-hour incubation at room temperature. Bound serum antibodies were then detected with a goat anti-mouse IgG-HRP secondary antibody (Southern Biotech 1030-05, Birmingham, AL) at 1:10,000 for 1 hour at room temperature. Lastly, plates were developed with TMB substrate for 5 minutes and stopped with 2 N sulfuric acid prior to measuring the absorbance at 450 nm on a BioTek plate reader. Titers were interpolated using non-linear regression in GraphPad Prism, with the cutoff value set at the limit of quantification for the assay (average absorbance of the blanks + 10 standard deviations). Statistical comparisons between groups were made using a Two-way ANOVA, while multiple corrections were made with the Tukey multiple comparisons test.

### B16F10 metastasis challenge model

On day 0, 250,000 B16F10 cells suspended in 100 μL of DPBS was administered into each C57BL/6 mouse intravenously via the tail vein using a heat lamp to facilitate the injections. Intraperitoneal administration of 200 μL of AvFc^ΔXF^, AvFc^Δlec^, or vehicle (AvFc formulation buffer, see above) at the indicated dose level began concurrently with tumor implantation and continued Q2D for a total of 6 doses (ending on day 10). Animals were monitored until day 21, at which point they were euthanized and the lungs removed for analysis. The tumor burden was calculated as the number of visible tumor nodules per lung per mouse. Statistical comparisons between treatment groups were made using the Kruskal-Wallis test, while multiple comparisons were made using Dunn’s test.

## Supporting information

SUPPORTING INFORMTION

## AUTHOR CONTRIBUTIONS

M.W.D., K.L.M., N.V.G. and N.M. wrote the manuscript. M.W.D. designed and performed protein expression and tumor challenge experiments. K.L.M. and N.V.G. assisted to perform parts of experiments. H.G. performed PET/CT analysis. H.K. performed glycan analysis. M.W.D., N.V.G., H.G., H.K., K.F. and N.M. analyzed data. N.M. conceived and supervised the study, and secured funding.

## ACKNOWLEDGMENTS

We thank Prof. Herta Steinkellner at Department of Applied Genetics and Cell Biology, University of Natural Resources and Life Sciences, Vienna, Austria, for providing ΔXylT/ΔFucT *Nicotiana benthamiana* seeds. This work was supported by an NIH grant to N.M.(R21-CA216447). M.D. was supported by an NIH Environmental Health Sciences Training Grant (T32-ES11564).

