## SUPPORTING INFORMTION for "Impact of glycoengineering and immunogenicity on the anti-cancer activity of a plant-made lectin-Fc fusion protein"

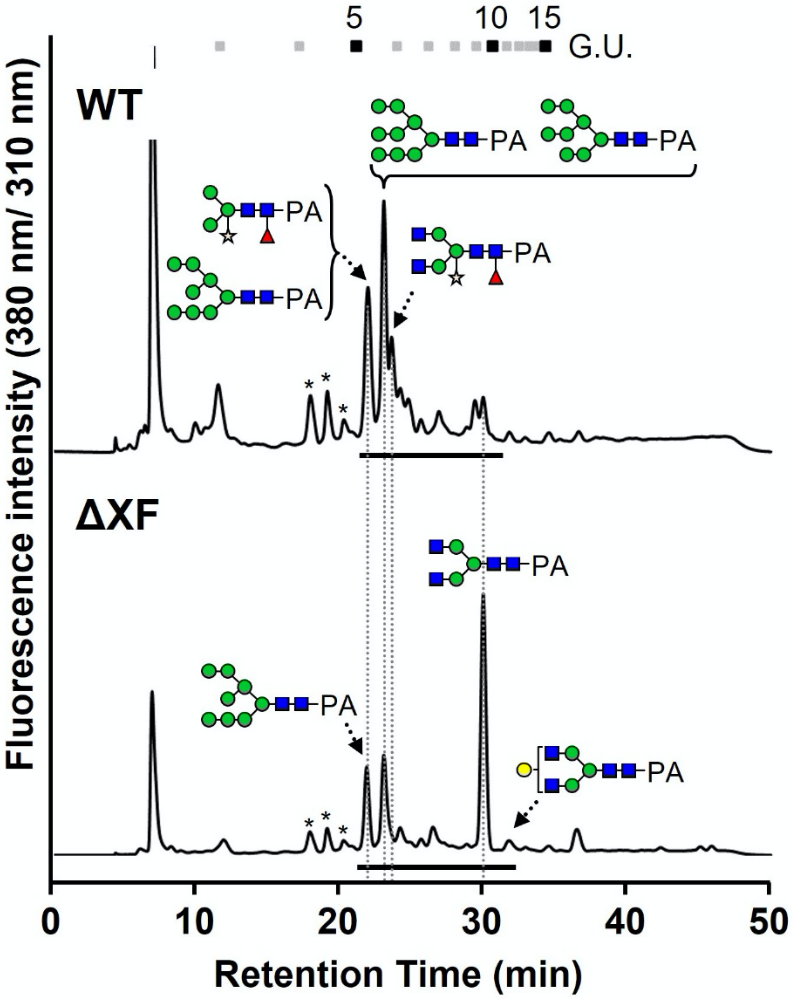


**Figure S1. Glycan profile of AvFc$^{WT}$ and AvFc$^{\Delta XF}$.**

Chromatograms show HPLC separation of PA-labeled glycan structures isolated from AvFc variants. Identification of Fc glycans by HPLC of WT and ΔXF AvFc shows the large presence of high-mannose glycans between both variants. WT AvFc also contains significant amounts of plant glycans containing α1,3-fucose and β1,2-xylose while ΔXF is devoid of them. Glycan symbols are drawn according to Symbol Nomenclature for Glycans (SNFG) nomenclature.

**A**

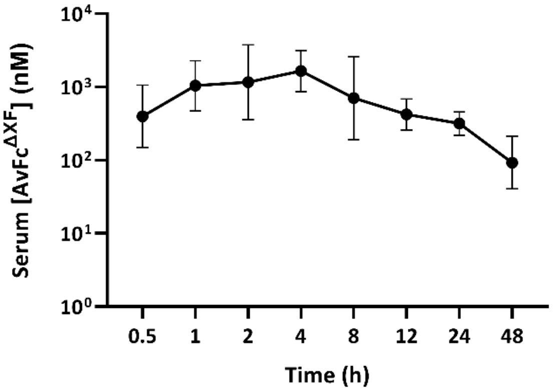


**B**

| Parameter | Unit | Value |
|---|---|---|
| Dose | ug | 500 |
| Lambda_z | 1/h | 0.037418 |
| $t_{1/2}$ | h | 18.5242 |
| Tmax | h | 4 |
| Cmax | nM | 1886.375 |
| AUCINF_obs | h*nM | 28226.57 |
| Vz_F_obs | ug/(nM) | 0.473398 |
| Cl_F_obs | ug/(h*nM) | 0.017714 |

**Figure S2. Pharmacokinetics of ΔXF AvFc in C57bl/6 mice.**

A pharmacokinetic profile for $AvFc^{\Delta XF}$ were measured in female C57bl/6 mice following a single intraperitoneal dose of 500 µg (25 mg/kg). Serum concentrations of $AvFc^{\Delta XF}$ were determined by gp120-binding ELISA at various time points and PK parameters were calculated using a non-compartmental model in Phoenix WinNonlin. The half-life of $AvFc^{\Delta XF}$ was determined to be approximately 18.5 hours.

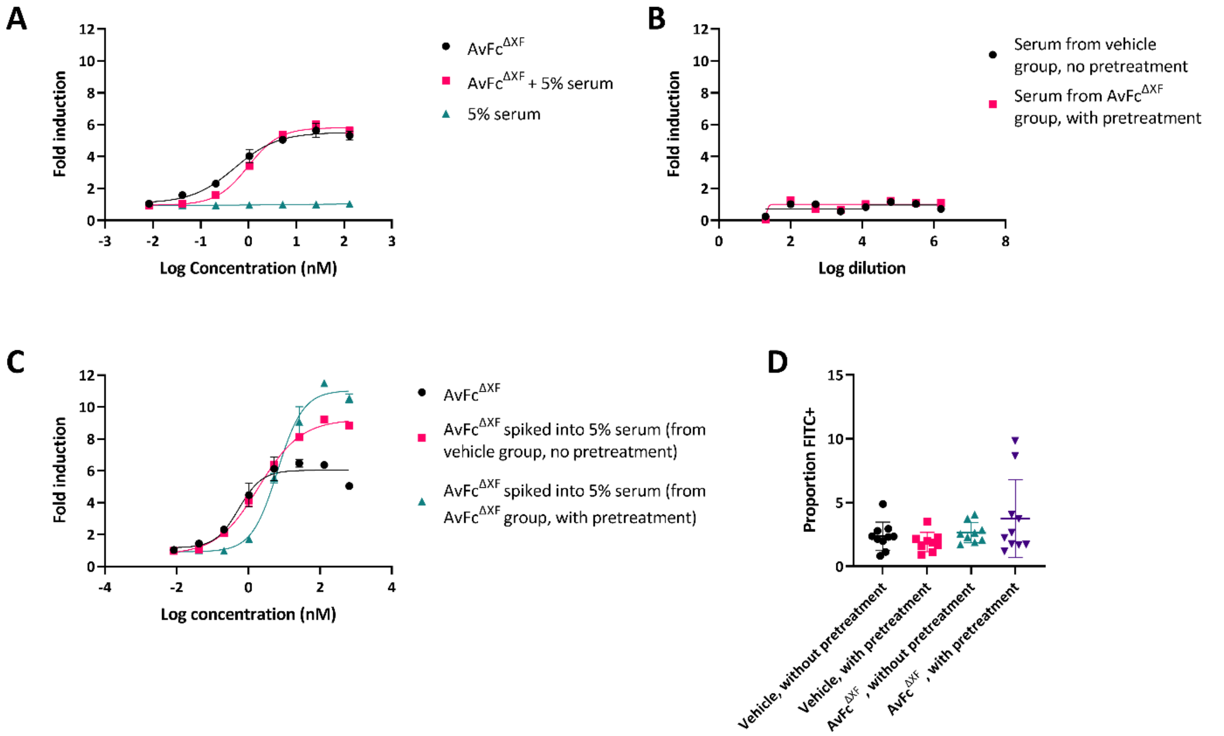


**Figure S3. Characterization of the impact of ADAs and ATAs.**

(A) ADCC reporter assay in the presence of mouse serum from before pretreatment. Serum was collected and pooled from mice prior to pretreatment with AvFc$^{\Delta XF}$, and the ADCC assay was performed as described previously with AvFc$^{\Delta XF}$ alone, AvFc$^{\Delta XF}$ spiked into a solution of 5% serum, and 5% serum alone. For all ADCC analyses, each data point represents the mean ± SD while the curves were fit with a 4-parameter non-linear regression model analysis in GraphPad Prism 9.2. No ADCC was induced by the 5% serum alone. Dose-response curves were nearly identical between AvFc$^{\Delta XF}$ alone and AvFc$^{\Delta XF}$ spiked into 5% serum, with some slight steepening of the curve. (B) ADCC reporter assay in the presence of terminal mouse serum alone. Serum was pooled from blood taken at euthanasia of each animal in each treatment group. Neither serum from animals in the non-pretreated, vehicle-treated group nor serum from the animals in the pretreated, AvFc$^{\Delta XF}$-treated group was capable of inducing ADCC on its own beginning at a 1:20 dilution. (C) ADCC reporter assay with AvFc$^{\Delta XF}$ spiked into terminal mouse serum. The ADCC assay was performed as in panel B with purified AvFc$^{\Delta XF}$ spiked into pooled serum from the non-pretreated, vehicle-treated group and the pretreated, $AvFc^{\Delta XF}$-treated group. Compared to $AvFc^{\Delta XF}$ alone, spiking into serum from the non-pretreated, vehicle-treated group resulted in an increase in the maximum fold induction from 6-fold to 9.2-fold and an increase in $EC_{50}$ from 0.53 nM to 1.94 nM. Spiking into serum from the pretreated, $AvFc^{\Delta XF}$-treated group resulted in an increase in the maximum fold induction from 6-fold to 11-fold and an increase in $EC_{50}$ from 0.53 nM to 6.44 nM. (D) Detection of anti-tumor antibodies with flow cytometry. Staining of B16F10 cells with a 1:10 dilution of pooled serum from each group followed by detection with a goat anti-mouse IgG FITC revealed no significant difference in the number of cells bound.

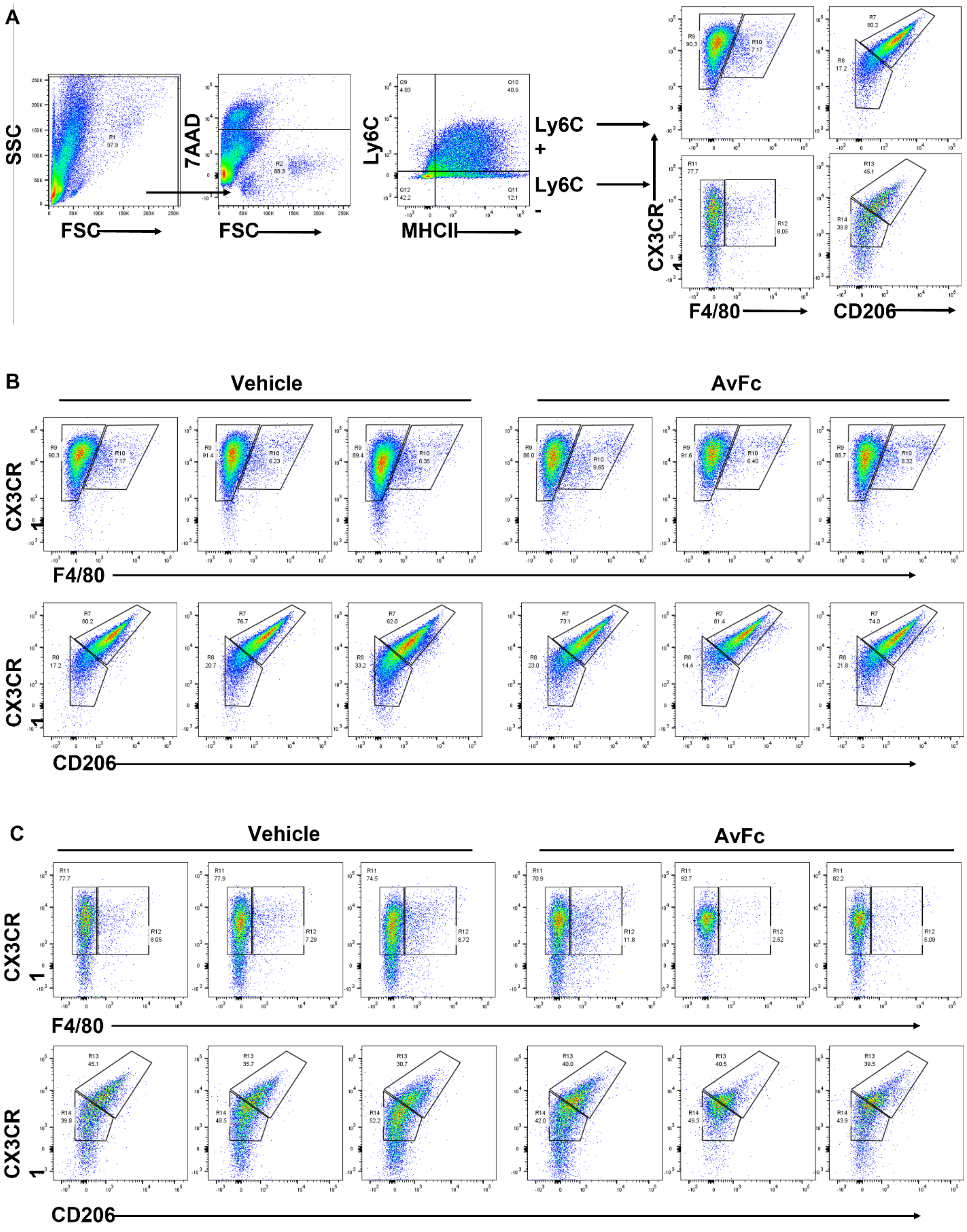
A
SSC
FSC
7AAD
FSC
Ly6C
MHCII
Ly6C +
Ly6C -
CX3CR1
F4/80
CD206
B
Vehicle
AvFc
CX3CR1
F4/80
CX3CR1
CD206
C
Vehicle
AvFc
CX3CR1
F4/80
CX3CR1
CD206

**Figure S4. The Fc-mediated effector functions of AvFc in B16F10 melanoma tumors involves the recruitment of non-classical monocytes.**

(A) Gating strategy used to analyze the infiltrating leukocytes into B16F10 melanoma tumors. Myeloid cells were gated from live cells by the expression of Ly6C and MHC class II (IA-IE), followed by the expression of $CX_3CR1$, F4-80 and CD206. (B) Proportions of classical monocytes defined as Ly6C+ myeloid cells gated from Vehicle and AvFc-treated mice. (C) Proportions non-classical monocytes defined as Ly6C- myeloid cells gated from Vehicle and AvFc-treated mice.

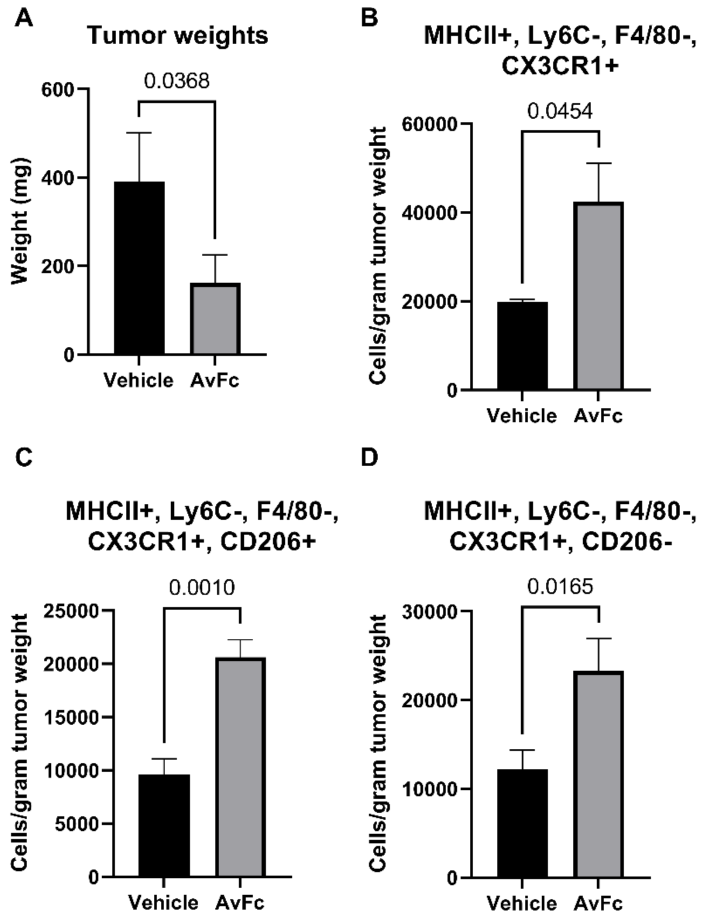


**Figure S5. Comparison of B16F10 tumor-infiltrating immune cells isolated from vehicle and AvFc-treated mice.**

Statistical comparisons of tumors weights and cell populations from Figure S4 were performed using Student's T test with Welch's correction in GraphPad Prism software. (A) Comparison of tumor weights at the time of removal from the animals shows a significant reduction in tumors from AvFc-treated animals (p=0.0368). (B-D) Treatment with AvFc was associated with a significant increase in the number of tumor-infiltrating non-classical monocytes (p=0.0454), and a significant increase in both the CD206+ and CD206- subpopulations of these cells (p=0.0010 and p=0.0165, respectively).

**Method S1. Immunophenotyping of B16F10 tumor-infiltrating immune cells**

B16F10 melanoma cells ($1x10^5$) were injected subcutaneously into the hind left flank of C57BL/6 (n=6/group) mice pre-treated with $AvFc^{\Delta XF}$ at 25 mg/kg or vehicle (AvFc formulation buffer). Tumor measurements were taken every day by using digital calipers until the tumor volume reached 500 $mm^3$, at this time the animals were euthanized, and the tumors dissected, weighed and minced for cell isolation. The minced cell suspension was digested in complete RPMI medium containing 2.5 mg/mL of Collagenase type IV (ThermoFisher Scientific) and 40 µg/mL of DNase I (MilliporeSigma, Saint Louis, MO) at 37 °C for 20 min under shaking conditions (200 rpm). Subsequently, the cells suspension was passed through a 40 µm cell strainer and the cell pellet resuspended and washed twice with FACS buffer, the cells were counted and incubated with 20 µg/mL of mouse gamma globulins to block FC-gamma receptors. A total of $1 \times 10^6$ Cells were stained for 30 min with 2 µg/mL of different combination of the following fluorochrome-labeled antibodies: anti-CD45eFluor450 or anti-CD45-FITC (30-F11), anti-CD3-FITC (17A2), anti-CD3-APC (17A2), anti-CD161 (NK1.1)-BV605 (PK136), anti-CD49b-PE (DX5), anti-CD107-AlexaFluor700 (1D4B), anti-CD335 (NKp46)-BV650 (29A1.4), anti-CD16.2-PE-Dazzle 594 (9E9), anti-CD11b-APC-Cy7 (M1/70), anti-CD11c-PE (N418), anti-IA-IE-BV421 (M5/114.15.5), anti-F4/80-PE-Cy7 (BM8), anti-Ly6G-APC (1A8), antiLy6C-AlexaFluor700 (HK1.4), anti-CX3CR1-BV605 (SA011F11), anti-CD206-BV650 (C068C2), anti-CD103-PE-Dazzle 594 (QA17A24), anti-CD80-BV605 (16-10A1), anti-CD69-BV650 (H1.2F3), anti-CD68-AlexaFluor700 (FA-11), anti-CDC86-PE-Dazzle 594 (GL-1), anti-CD4-BV605 (RM4-5), anti-CD8-BV650 (53-6.7), anti-IFNγRβ-APC (MOB-47), anti-CD69-FITC (H1.2F3), anti-IL-33R-PE-Dazzle 594 (DIH4), anti-CD62L-APC-Cy7 (MEL-14), anti-TCRβ-PE-Cy7 (H57-597), anti-IL23R-BV421 (12B2B64) and anti-TCRγ/δ-PE (UC7-13D5). After two washing steps the cells were incubated with 7-aminoactinomycin D for 15 minutes and analyzed with a BD LSRFortessa$^{TM}$ flow cytometer and the data processed with FlowJo_v10.8.0_CL software.
